# Response of the multiple-demand network during simple stimulus discriminations

**DOI:** 10.1101/296509

**Authors:** Tanya Wen, Daniel J Mitchell, John Duncan

## Abstract

The multiple-demand (MD) network is sensitive to many aspects of task difficulty, including such factors as rule complexity, memory load, attentional switching and inhibition. Many accounts link MD activity to top-down task control, raising the question of response when performance is limited by the quality of sensory input, and indeed, some prior results suggest little effect of sensory manipulations. Here we examined judgments of motion direction, manipulating difficulty by either motion coherence or salience of irrelevant dots. We manipulated each difficulty type across six levels, from very easy to very hard, and additionally manipulated whether difficulty level was blocked, and thus known in advance, or randomized. Despite the very large manipulations employed, difficulty had little effect on MD activity, especially for the coherence manipulation. Contrasting with these small or absent effects, we observed the usual increase of MD activity with increased rule complexity. We suggest that, for simple sensory discriminations, it may be impossible to compensate for reduced stimulus information by increased top-down control.

## 1. Introduction

Diverse studies examining a range of cognitive demands have found of a set of frontal-parietal regions that are consistently involved in a variety of tasks, ranging from response inhibition to working memory to decision making (e.g., Duncan & Owen, 2000; Fedorenko, Duncan, & Kanwisher, 2013; Niendam et al., 2012; Stiers, Mennes, & Sunaert, 2010). Included in this pattern are regions of the dorsolateral prefrontal cortex, extending along the inferior/middle frontal gyrus (IFG/MFG), and including a posterior-dorsal region close to the frontal eye field (pdLFC), parts of the anterior insular cortex (AI), pre-supplementary motor area and adjacent anterior cingulate cortex (pre-SMA/ACC), and intraparietal sulcus (IPS). Together they have been termed the multiple demand (MD) network (Duncan, 2010), cognitive control network (Niendam et al., 2012), or task positive network (Fox et al., 2005).

Activity in the MD network increases with increases in many kinds of task difficulty or demand, such as with additional subgoals (e.g., Farooqui et al., 2012), greater working memory demand (Dara et al., 1997), resisting strong competitors (e.g., Baldauf & Desimone 2014), task switching (e.g., Wager et al., 2004), or a wide range of other task demands (e.g., Crittenden & Duncan, 2014; Jovicich et al., 2001; Marois, Chun, & Gore, 2004; Woolgar, Afshar, Williams, & Rich, 2015). Increased activity in more difficult conditions can also be accompanied by stronger information coding, shown by multivoxel pattern analysis (e.g., Woolgar, Afshar, et al., 2015; Woolgar, Hampshire, Thompson, & Duncan, 2011; Woolgar, Williams, & Rich, 2015). Reflecting these widespread effects of demand, the MD network has been suggested to implement top-down attentional control, optimally focusing processing for the requirements of a current task (Miller & Cohen, 2001; Duncan, 2013; see also Norman & Shallice, 1980).

One simple way to manipulate task difficulty is through the quality of stimulus information. Some experiments have shown clear MD responses as stimulus discriminability decreases (e.g., Crittenden et al., 2014; Deary et al., 2004; Holcomb et al., 1998; Jiang & Kanwisher, 2003; Sunaert, Van Hecke, Marchal, & Orban, 2000; Woolgar et al., 2011), but this has not always been the case (Cusack, Mitchell, & Duncan, 2010; Dubis et al., 2016; Han & Marois, 2013; Muller-Gass & Schroger, 2007). For example, Cusack et al. (2010) contrasted hard and easy trials of a task in which participants had to detect a barely perceptible ripple in an oscillating dot field and found no neural activation differences between the two sensory difficulty levels, despite substantial differences in behavioral performance, and robust BOLD contrast to a different task manipulation (attention switching).

In an important study, Han and Marois (2013) investigated activity in parts of the MD system during a task in which three letter targets were to be identified in a rapid stream of digit nontargets. In the baseline condition, the three letters occurred in immediate succession; to increase demand, they either inserted a nontarget into the series of three targets, or reduced exposure duration. While activity in frontal-parietal areas increased with the addition of a distractor, exposure duration had little effect. To interpret their findings, Han and Marois (2013) appealed to the distinction made by Norman and Bobrow (1975), between data-limited and resource-limited behavior. Norman and Bobrow (1975) proposed that, for any task, some function (the performance-resource function or PRF) relates performance to investment of attentional resources. When this function is increasing, behavior is said to be resource-limited, and additional investment is repaid by improved performance. When the function asymptotes, further investment has no positive effect, and performance is said to be data-limited. In line with a link of MD activity to attentional investment, Han and Marois (2013) used these ideas of data- and resource-limitation to explain their findings. They proposed that, in their task, brief exposure duration created data limits, which could not be offset by increased fronto-parietal recruitment, while adding a distractor introduced resource limits by calling for increased attentional focus.

In general it is not known when performance will be resource- or data-limited, but within this general framework, many patterns of results are possible. Figure 1A illustrates a case in which, as difficulty level varies, there is no reason to expect increased attentional allocation. In this case, PRFs asymptote at different performance levels for the different levels of task difficulty, but across difficulty levels, the asymptote occurs at the same level of allocated resource. Figure 1B illustrates an opposite case, with increased task difficulty potentially offset by increased resource allocation. This uncertainty over the role of attentional investment in different cases of perceptual discrimination could help to explain disparate results in the literature, with some cases (e.g. Han and Marois (2013), manipulation of exposure duration) more resembling Figure 1A, and others (e.g. Han and Marois (2013), distractor manipulation) more resembling Figure 1B.

**Figure 1.**
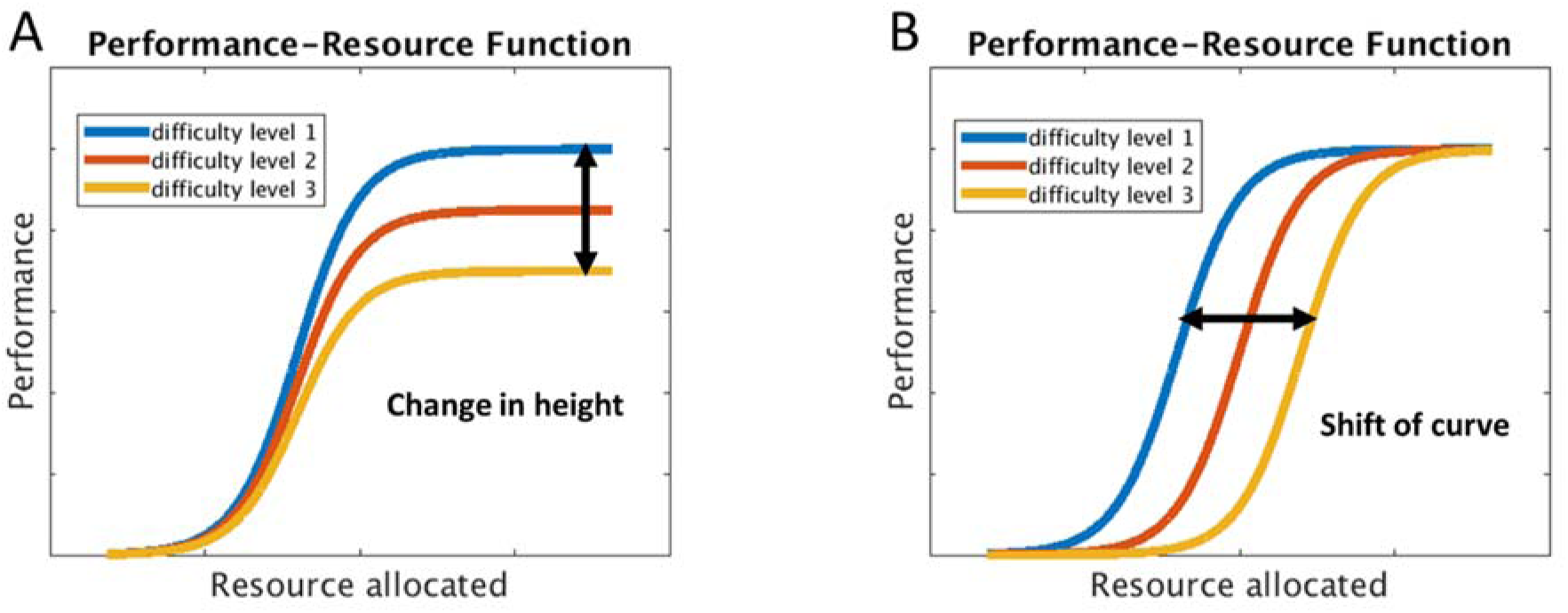
Theoretical performance-resource function (PRF) plots. (A). Difficulty might change the asymptote of of the PRF. Increased difficulty cannot be offset by increased resource allocation. (B). Difficulty might shift the PRF. Increased difficulty can be offset by increased resource allocation.

In our first experiment, we sought to strengthen the evidence that, for simple sensory discriminations, MD activity can be rather independent of task difficulty, providing an exception to the “multiple demand” pattern. For this purpose we used a motion discrimination task with two kinds of difficulty manipulation – motion coherence and salience of task-irrelevant dots. For the strongest possible effect, we manipulated both variables over a wide range, moving performance from close to ceiling to close to chance. In the task demand literature, several studies have shown that, as opposed to a monotonic increase of MD activity with task difficulty, there was an inverted U-shape response (Callicott et al., 1999; Linden et al., 2003), or a plateau after a certain difficulty level (Marois & Ivanoff, 2005; Todd & Marois, 2004; Mitchell & Cusack, 2008). A possible interpretation is that MD activity initially increases with task demands, but plateaus or even declines once the task becomes impossible even with maximal attention. In our study we examined MD activity over the full range of possible task difficulties.

In addition to manipulating both aspects of difficulty over a wide range, between participants we varied whether difficulty levels were mixed or blocked. In the mixed design, levels of difficulty were presented in random order, without advance cueing of the level to be experienced on a given trial. In contrast, difficulty level was known in advance in the blocked design. With this manipulation, we asked whether MD activity is driven more proactively, by expectancy of forthcoming demand, or more reactively, when high demand is experienced on a current trial.

Finally, in modelling our fMRI data, we attempted to remove effects of decision time, expected to increase with either sensory or selection difficulty. In two prior studies of motion coherence, trials were modelled simply as events, without regard for their duration (Kayser et al., 2010a, 2010b). In this case, greater brain activity associated with decreasing motion coherence may simply have reflected longer processing times. To diminish such effects, our fMRI model explicitly included decision time for each trial.

Though PRF shapes are generally unknown, our use of two different demand manipulations afforded the possibility of different outcomes. In particular, we expected that top-down control could be especially important in the irrelevant-dots condition, leading to larger effects of demand on MD activity. Though Experiment 1 showed results in line with this expectation, they occurred against a background of generally weak effects, and no significant difference between the two manipulations. In Experiment 2 we reexamined coherence and irrelevant-dots conditions in a new group of participants, and compared these sensory demands with a manipulation of rule complexity.

## 2. Methods

### 2.1 Experiment 1

#### 2.1.1 Participants

Participants were randomly assigned to either the blocked or mixed group, with this variable manipulated between participants to minimize carryover effects. A total of 40 participants took part in the experiment. Twenty-one participants (9 male, 12 female, ages 19-31, mean = 25.7) took part in the blocked group, and nineteen participants (11 male, 9 female, ages 19-36, mean = 23.9) took part in the mixed group. Participants were recruited from the volunteer panel of the MRC Cognition and Brain Sciences Unit and paid to take part. An additional 16 participants were excluded (10 participants had excessive motion > 5mm, and another 6 had poor performance with accuracies more than three median absolute deviations below the median in at least one condition). All participants were neurologically healthy, right-handed, with normal hearing and normal or corrected-to-normal vision. Procedures were carried out in accordance with ethical approval obtained from the Cambridge Psychology Research Ethics Committee, and participants provided written, informed consent before the start of the experiment.

#### 2.1.2 Experimental Setting and Design

Each participant performed two conditions of a motion coherence task, referred to here as the coherence condition and the irrelevant-dots condition. Each condition spanned six levels of difficulty. Difficulty type and level served as within-subject factors. This resulted in a group (blocked vs. mixed) × difficulty type (coherence vs. irrelevant-dots) × difficulty level (level 1 ~ level 6) design.

#### 2.1.3 Stimuli and Procedures

The task structure was similar for both blocked and mixed designs (see Figure 2). On each trial, participants were presented with a random dot kinematogram (RDK) displayed for 200 ms, with an interval of 2-3 s between RDKs of successive trials. Participants were asked to judge the direction of the dominant dot motion, leftward or rightward. They were given 2 s to press one of two response buttons (up or down) to indicate their decision. The mapping between stimulus (left or right) and response (up or down) varied randomly between blocks, ensuring that, across the whole experiment, there were equal numbers of left-up/right-down and left-down/right-up trials for each difficulty level in each condition.

**Figure 2.**
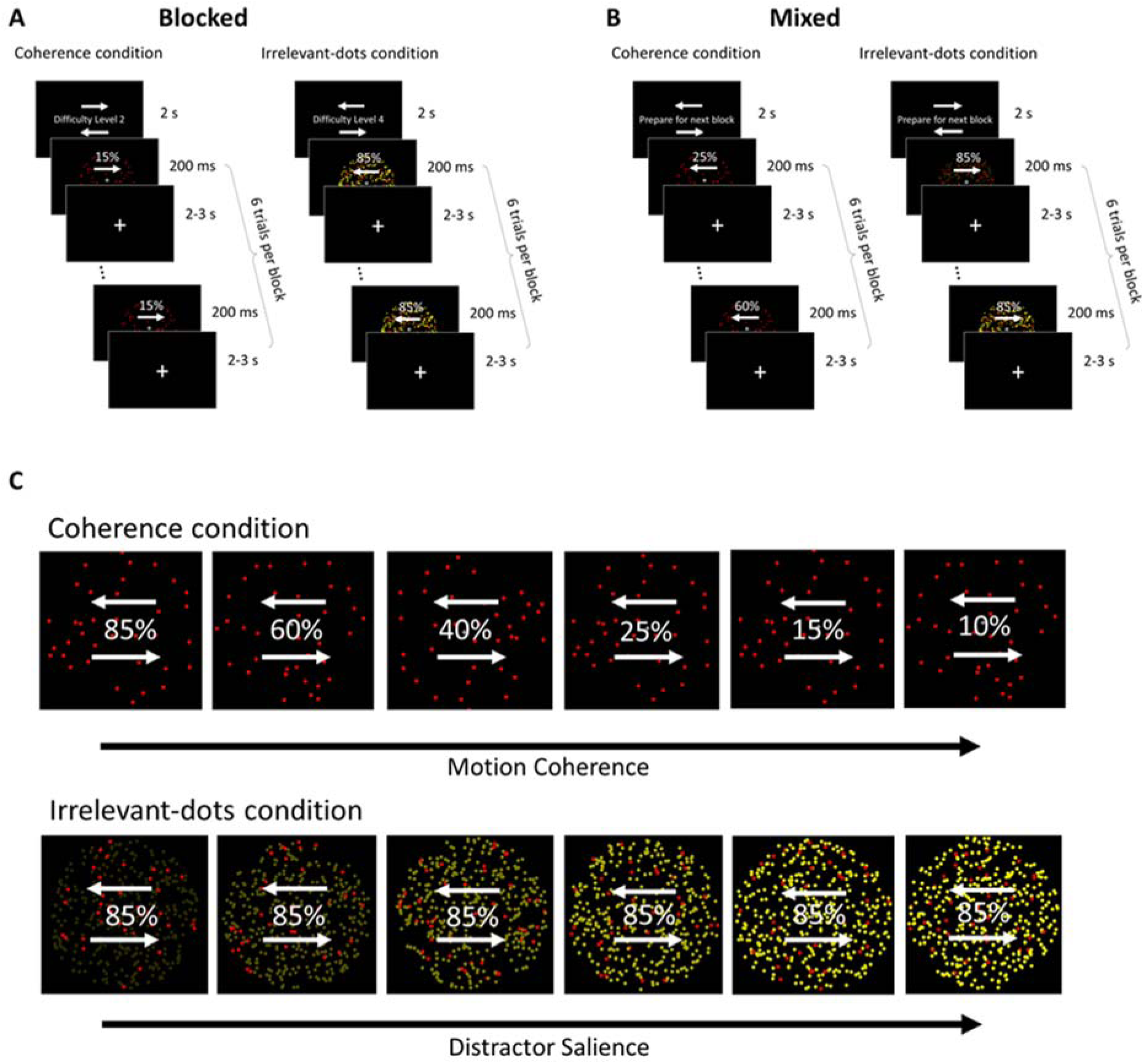
An example of one block of each condition in the blocked design (A) and the mixed design (B). At the beginning of each block, participants were given the response mappings and were cued to prepare for that block. At the beginning of each trial, a 200 ms RDK appeared, followed by a 2-3 s fixation period, during which participants were to judge the motion direction of the red dot field. The percentages indicate degree of motion coherence and were not shown on the actual display. (C) An illustration of the 6 levels of the coherence and irrelevant-dots conditions. In the coherence condition, difficulty was manipulated by decreasing the motion coherence of the dot field. In the irrelevant-dots condition, difficulty was manipulated by increasing the salience of the yellow dots. The percentages indicate the degree of motion coherence and were not shown on the actual display.

Trials were run in blocks of six. At the beginning of each block, the response mappings were displayed on the screen during a 2 s instruction period preceding the first trial, and remained on the screen throughout the entire block. The response mappings were indicated by two arrows above and below each other within the dot field aperture, with one arrow pointing right and the other pointing left. In the blocked version, information about the difficulty level of the following block was also shown on the screen (e.g., ‘Difficulty Level is 1’) during the instruction period; while in the mixed version, participants were shown the instructions ‘Prepare for Next Block’. The difficulty level of the six trials in a block remained the same in the blocked design, covering all six difficulty levels every six blocks; while in the mixed design each block contained one trial of each difficulty level randomized within the block.

Participants were given two practice blocks of each condition (coherence and irrelevant-dots) before entering the scanner. Within the scanner, each condition constituted a separate run, each lasting ~18 minutes. The order of the two conditions was counterbalanced between participants. In each run there were 60 blocks in total, thus 60 trials per difficulty level, and 30 trials of each left/right motion direction in each difficulty level.

Figure 2C illustrates coherence and irrelevant-dots conditions across the six difficulty levels. In each RDK there were 64 red dots (RGB channels [112.5, 0, 0]) moving dominantly either left or right for 200 ms (circular aperture with diameter of visual angle ~8.5°, dot size = 12 pixels diameter, dot speed = 5 pixels/s, dot lifetime = 5 frames). In the coherence condition, only red dots were present, and difficulty was manipulated by decreasing the percentage of the dots that were moving coherently. The six difficulty levels corresponded to 85%, 60%, 40%, 25%, 15%, or 10% of the dots moving either left or right, while the remaining dots moved in random directions. In the irrelevant-dots condition, the red dots were fixed at 85% coherence, but an additional distractor dot field was present. The distractor dot field consisted of 576 yellow dots, all of which moved randomly, with a net direction of zero (dot size = 12 pixels, dot speed = 7 pixels/s, dot lifetime = 5 frames). Participants were asked to ignore the yellow dots, and judge the dominant motion direction of the red dots. Six levels of difficulty were created by increasing the salience of the yellow distractors (RGB channels [21.25, 21.25, 0], [42.5, 42.5, 0], [85, 85, 0], [127.5, 127.5, 0], [191.25, 191.25, 0], and [255, 255, 0]). These values were selected from previous pilot testing.

Visual stimuli were displayed on a 1920×1080 (visual angle 25.16×14.31°) screen with a refresh rate of 60 Hz, which the participants viewed through a mirror placed on top of the head coil. The distance between the participant and screen was approximately 1565 mm. Stimulus presentation was controlled using the Psychophysics Toolbox (Brainard, 1997) in Matlab (2014a, Mathworks, Natick, WA).

#### 2.1.4 Imaging Parameters and Data Analyses

Scanning took place on a 3T Siemens Tim Trio scanner. Functional images were acquired using a multi-band gradient-echo echo-planar imaging (EPI) pulse sequence (TR = 1100 ms, TE = 30 ms, flip angle = 62°, 78 × 78 matrices, slice thickness = 2.5 mm, no gap, voxel size 2.5 mm × 2.5 mm × 2.5 mm, 48 axial slices covering the entire brain, 3 slices acquired at once). The first 10 volumes served as dummy scans and were discarded to avoid T1 equilibrium effects. Field maps were collected at the end of the experiment (TR = 400 ms, TE = 5.19 ms / 7.65 ms, flip angle = 60°, 64 × 64 matrices, slice thickness = 3 mm, 25% gap, resolution 3 mm isotropic, 32 axial slices). High-resolution anatomical T1-weighted images were acquired for each participant using a 3D MPRAGE sequence (192 axial slices, TR = 2250 ms, TI = 900 ms, TE = 2.99 ms, flip angle = 9°, field of view = 256 mm × 240 mm × 160 mm, matrix dimensions = 256 × 240 × 160, 1 mm isotropic resolution).

The data were preprocessed and analyzed using the automatic analysis (aa) pipelines and modules (Cusack et al., 2014), which called relevant functions from Statistical Parametric Mapping software (SPM 12, http://www.fil.ion.ucl.ac.uk/spm) implemented in Matlab (The MathWorks, Inc., Natick, MA, USA). EPI images were realigned to correct for head motion using rigid-body transformation, unwarped based on the field maps to correct for voxel displacement due to magnetic-field inhomogeneity, and slice time corrected. The T1 image was coregisted to the mean EPI, and then coregistered and normalized to the MNI template. The normalization parameters of the T1 image were applied to all functional volumes. The functional data were high-pass filtered with a cutoff at 1/128 Hz, and spatial smoothing of 10 mm FWHM was applied.

Statistical analyses were performed first at the individual level using general linear modeling (GLM). For correct trials, separate regressors were created for each combination of condition and difficulty level. As errors can be a strong driver of MD activity (Kiehl et al., 2000; Ullsperger & Cramon, 2004), error trials and no-response trials were removed using separate regressors. A separate regressor was created for the cue period at the start of each block. All regressors were created by convolving the interval between stimulus onset and response with the canonical hemodynamic response function.

Frontoparietal MD ROIs were taken from Fedorenko et al. (2013) (http://imaging.mrc-cbu.cam.ac.uk/imaging/MDsystem). These included: anterior cingulate and pre-supplementary motor area (ACC/pre-SMA), anterior insula (AI), posterior-dorsal prefrontal cortex (pdLFC), intraparietal sulcus (IPS), and three foci along the middle frontal gyrus (anterior, middle, and posterior MFG). Beta estimates for correct trials were averaged within each ROI and analyzed using a 4-way mixed ANOVA, with factors group (blocked vs. mixed), difficulty type (coherence vs. irrelevant-dots), difficulty level (level 1 ~ level 6), and ROI (7 MD ROIs). We also performed the same ANOVA on the motion-sensitive visual area MT, using the ROI defined in the SPM anatomy toolbox.

To examine whether individuals showed significant linear effects for difficulty level, we performed a GLM on average activity across the entire MD network, using the MarsBaR toolbox (Brett et al., 2002). Positive linear contrasts (activity increasing with task difficulty) were defined as [-5 -3 -1 +1 +3 +5], across difficulty levels 1 to 6, and negative linear contrasts as [+5 +3 +1 -1 -3 -5].

An additional whole-brain voxelwise analysis was also performed, to detect any regions that showed a significant positive linear trend for difficulty level, separately for each difficulty type. Activation maps were thresholded at p < 0.05 (FDR corrected).

### 2.2 Experiment 2

A separate set of participants (n = 24, 18 female, ages 19-30, mean = 24.4) was recruited to perform the coherence and irrelevant-dots conditions along with an additional rule condition. 5 additional participants were excluded (1 because the top of the brain was not acquired, 2 who had head movements > 5 mm, and 2 who had outlier reaction times more than three median absolute deviations above the median). All participants performed the blocked design.

The coherence and irrelevant-dots conditions used a subset (levels 1, 3, and 5) of the same stimuli as previously described, except this time (Figure 3A) the dots went in one of four directions (15°, 65°, 115°, 165°). The stimuli and response-mappings for the rule condition are illustrated in Figure 3. Participants responded using the index and middle finger of each hand on the four buttons of a response pad (left hand middle finger for the first button, left hand index finger for the second button, right hand index finger for the third button, and right hand middle finger for the fourth button), with a direct spatial mapping between stimulus direction and response key (Figure 3B, level 1. The dot fields in the rule difficulty condition had high coherence (85%) and did not include yellow dots. However, rule complexity was manipulated using three different mappings between stimulus direction and response key (Figure 3B).

**Figure 3.**
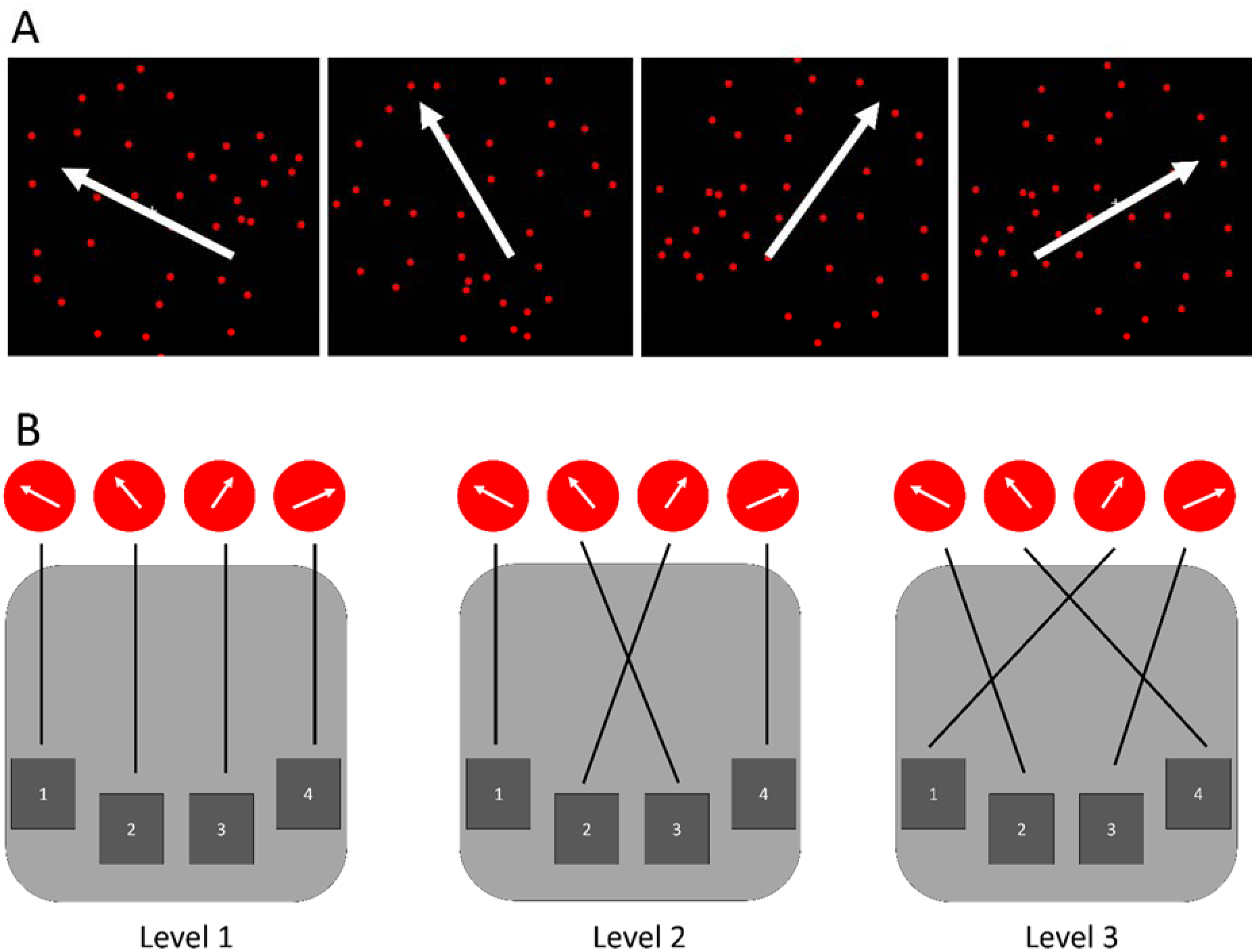
(A) Stimuli in Experiment 2. Red dots moved in one of four possible directions (15°, 65°, 115°, 165°). Additional yellow dots (not shown) were added only in the irrelevant-dots condition. (B) Participants were asked to use a response pad to indicate the direction of the moving dots by pressing the corresponding button. A direct mapping (level 1) was used for coherence and irrelevant-dots conditions, while all 3 rules were used in the rule difficulty condition.

At the beginning of each block, participants were presented with a cue indicating the difficulty level of the block. After processing the cue, they were able to press a button to begin a consecutive 6 trials of that condition. Each RDK was presented for 200 ms. Participants had up to 10 s to respond, and after a button was pressed, a 2 s ISI preceded the next trial. At the cue for the next block, participants were given feedback of their performance accuracy as well as mean reaction time for the previous block.

Participants were given two practice blocks of each condition (coherence, irrelevant-dots, and rule) before entering the scanner. Within the scanner, each condition constituted a separate run. The order of the three conditions was counterbalanced across participants. In each run there were 24 blocks in total, thus 48 trials per difficulty level.

## 3. Results

### 3.1 Experiment 1

#### 3.1.1 Behavioral results

As shown in Figure 4A, accuracy decreased while reaction times increased with difficulty in both groups. Overall, proportion correct decreased from a mean of 93.5%, to a mean of 63.0% from the easiest to the hardest difficulty level.

**Figure 4.**
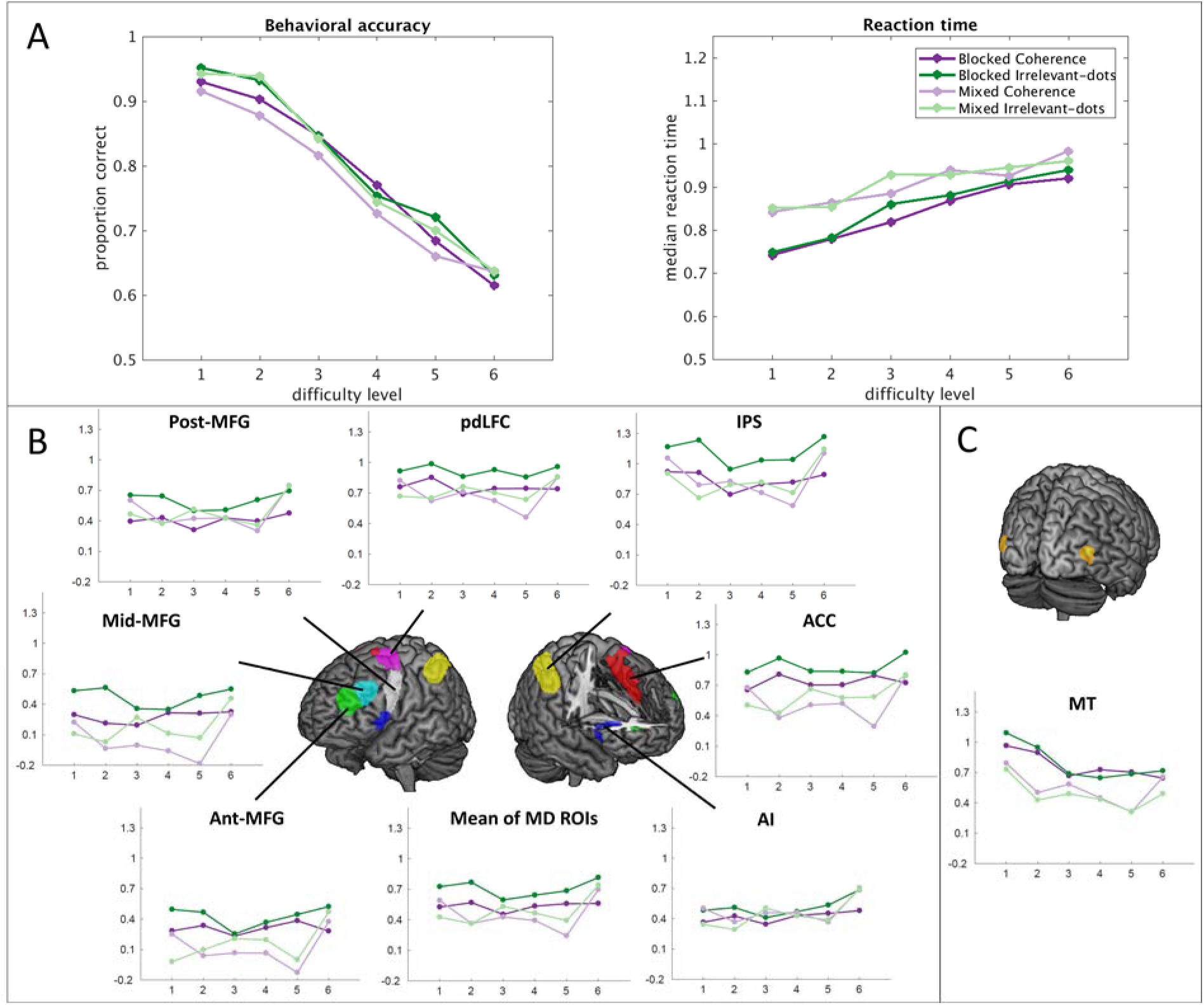
Experiment 1. (A). Behavioral results for each group and condition. Left: accuracy; right: reaction time). (B). ROI results of MD regions. (C). ROI results of MT. Graphs plot the beta values for each condition averaged across bilateral ROIs. Error bars represent the standard error of the mean.

3-way ANOVAs of group (blocked vs. mixed) × condition (coherence vs. irrelevant-dots) × difficulty level (level 1 ~ level 6) were performed on proportion correct and reaction time. The Greenhouse-Geisser correction was used to correct for non-sphericity. For accuracy, there were significant main effects of condition, F(1,38) = 4.1, p = 0.049, with slightly higher accuracy for irrelevant-dots, and difficulty level, F(5,190) = 240.36, p < 0.001. There was no main effect of group, F(1,38) = 0.3, p = 0.61, and no interactions. Analysis of reaction times showed a significant main effect of difficulty level, F(5,190) = 69.6, p < 0.001. The reaction time analysis also showed a significant but small interaction of condition × difficulty level, F(5,190) = 3.8, p < 0.01, but no other effects.

#### 3.1.2 fMRI results

Results averaged over bilateral MD regions are shown in Figure 4B, separately for each condition and group. As a first step, we used 2-way ANOVAs (group × difficulty level) to examine the effect of difficulty separately in each condition, averaged across all MD ROIs. In the coherence condition, the effect of difficulty was not close to significance, F(5,190) = 1.7, p = 0.14. For the irrelevant-dots condition, in contrast, increased activity across difficulty levels was significant, F(5,190) = 3.1, p = 0.01. There were no significant interactions with group.

Next we tested linear increases with difficulty level in each condition separately. For the coherence condition, there were no linear effects of difficulty in the blocked group, F(1,20) = 0.1, p = 0.73, or in the mixed group, F(1,18) = 0.04, p = 0.84. For the irrelevant-dots condition, there was a significant linear trend in the mixed group, F(1,18) = 5.6, p = 0.03, but no effects in the blocked group, F(1,20) = 0.3, p = 0.61.

Next, to compare across groups, conditions and ROIs, we used 4-way ANOVA with factors group (blocked vs. mixed), condition (coherence vs. irrelevant-dots), difficulty level (level 1 – level 6), and ROI (7 MD ROIs, averaged across the two hemispheres). This analysis revealed a significant main effect of difficulty level, F(5,190) = 4.1, p < 0.05. Tests of a linear contrasts across difficulty levels showed a significant positive trend, F(1,38) = 4.3, p = 0.04. Critically, there was no significant interaction of condition and difficulty level, F(5,190) = 0.3, p = 0.89, though a significant interaction between group and difficulty level, F(5,190) = 2.7, p = 0.03 reflected somewhat different activity profiles for blocked and mixed participants (see Figure 4). Though absolute activation levels differed over MD ROIs, trends were largely similar across ROIs (Figure 4B). The ANOVA showed a significant main effect of ROI, F(6,228) = 37.3, p < 0.001, along with a significant interaction of ROI and difficulty level, F(30,1140) = 2.6, p = < 0.001.

To check that our ROI analysis did not miss important effects elsewhere in the brain, we also carried out a standard whole-brain analysis, combining data for blocked and mixed groups, and testing for a linear increase across difficulty levels (see Methods). For the coherence condition, no significant voxels were found anywhere in the brain. For the irrelevant-dots condition, beyond the expected large increases in visual cortex, the test showed scattered, small regions close to our MD ROIs, including preSMA/ACC, AI, and regions of lateral frontal cortex.

Lastly, we tested for significant linear increases or decreases in individual participants (see Methods), again combining blocked and mixed groups, and using a whole-MD ROI. With 40 participants and an alpha level of .05, for each test we should expect 2 participants to be judged significant by chance. For the coherence condition, there were 10 significant increases but also 8 significant decreases. For the irrelevant-dots condition, there were 17 significant increases and 3 significant decreases. Overall, these results are broadly similar to those from standard random-effects analysis, but further, they suggest a significant degree of heterogeneity between participants.

ROI results for MT are shown in Figure 4C. In line with prior results (Rees, Friston, & Koch, 2000), MT activity generally declined with increasing difficulty of extracting the motion signal. A 3-way ANOVA with factors group (blocked vs. mixed), condition (coherence vs. irrelevant-dots), and difficulty level (level 1 – level 6) showed a significant main effect of difficulty level, F(5,190) = 11.7, p < 0.001. There was also a group by level interaction, F(5,190) = 2.9, p = 0.03, reflecting a stronger difficulty effect in the blocked group. No other significant effects were observed. Tests of within-subjects contrasts on difficulty level showed a significant negative linear trend, F(1,38) = 58.6, p < 0.001.

### 3.2 Experiment 2

#### 3.2.1 Behavioral results

In Experiment 2, participants performed the coherence, irrelevant-dots, and rule conditions. Behavioral data are shown in Figure 5A. Separate condition (sensory, selection, and rule) × difficulty level (level 1 ~ level 3) ANOVAs were performed on accuracy and reaction time. For accuracy, there were main effects of difficulty level, F(2,26) = 187.2, p < 0.001, and condition, F(2,46) = 64.2, p < 0.001. There was a significant interaction between condition and difficulty level, F(4,92) = 69.2, p < 0.001, reflecting generally high accuracy across difficulty levels in the rule condition. Reaction times also showed significant main effects of difficulty level, F(2,46) = 112.3, p < 0.001, and condition, F(2,46) = 7.0, p < 0.01, and a significant interaction between condition and difficulty level, F(4,92) = 10.6, p < 0.001.

**Figure 5.**
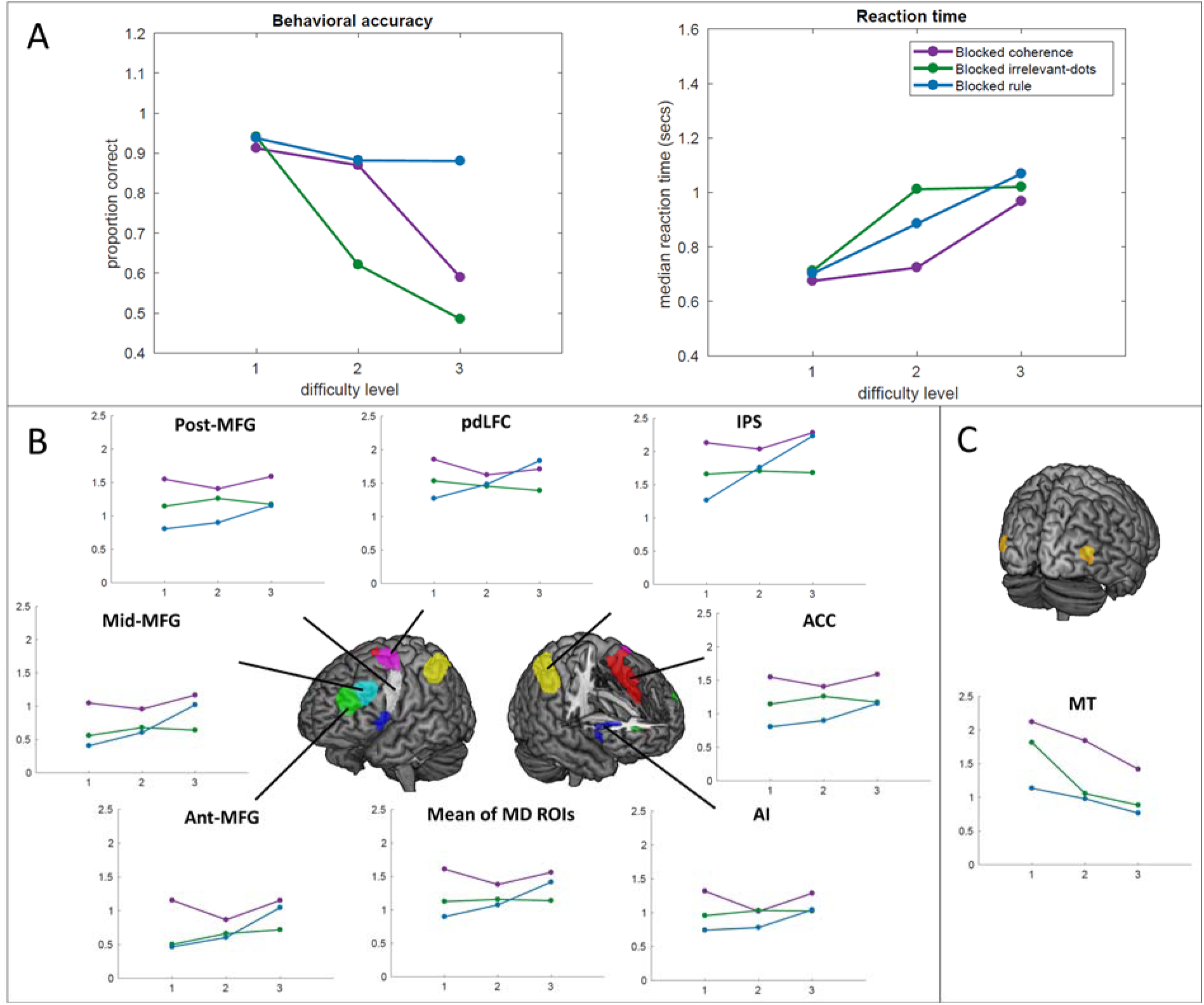
Experiment 2. A. Behavioral results each group and condition. Left: accuracy; right: reaction time. B. ROI results of MD regions. C. ROI results of MT. Graphs plot the beta values for each condition averaged across bilateral ROIs. Error bars represent the standard error of the mean.

#### 3.2.2 fMRI results

Results for MD regions are shown in Figure 5B. In this experiment, neither coherence nor irrelevant-dots conditions showed any trend towards increasing activity with increasing difficulty, in contrast to results from the rule condition. In the MD regions, we found significant 2-way interactions of condition and difficulty level, F(4,92) = 2.9, p < .05, and ROI and difficulty level, F(12,276) = 3.6, p < .01, as well as a 3-way interaction of condition, difficulty level, and ROI, F(24,552) = 2.2, p < .05. Further we tested for linear increases across difficulty levels in each condition separately. Difficulty level showed a significant linear trend in the rule condition, F(1,23) = 4.5, p < .05; however, there were no linear trends for either coherence, F(1,23) = 1.1, p = 0.32 or irrelevant -dots F(1,23) = 0.1, p = 0.76.

Whole-brain analysis showed no voxels with a significant linear increase across difficulty levels, either for coherence or irrelevant-dots conditions. For the rule condition, significant effects were found in lateral parietal and lateral frontal cortex bilaterally. Tests on individual participants showed 9/24 increases and 6/24 decreases in the coherence condition; the same in the irrelevant-dots condition; and in the rule condition, 14/24 increases and 2/24 decreases.

Results for MT are shown in Figure 5C. There were significant main effects of difficulty level, F(2,46) = 17.1, p < 0.001, and condition, F(2, 46) = 5.6, p = 0.01, but no interactions.

## 4 Discussion

The characteristic of fronto-parietal “multiple-demand” regions is increased activity for many different kinds of task difficulty. Here, we pursued mixed previous findings suggesting a partial exception – in some cases, MD activity seems largely insensitive to the difficulty of sensory discriminations. To obtain the strongest possible test of such insensitivity, using a motion coherence task, we manipulated two aspects of sensory difficulty over the widest possible range, from very easy to close to chance. We also compared results with difficulty fixed or variable across a block of trials. To ensure good power, across two experiments we tested a total of 64 participants.

Clear results were obtained for a manipulation of motion coherence. Across experiments and mixed or blocked variations of difficulty level, there was no hint of consistently increased MD activity as performance changed from close-to-ceiling to close-to-chance. Following Han and Marois (2013), these results can be interpreted in terms of the distinction drawn by Norman and Bobrow (1975) between data and resource limitations. For motion coherence, the results suggest a scenario similar to that depicted in Figure 1A: decreasing coherence simply decreases the quality of sensory data, and this cannot be offset by increased attentional focus or top-down control.

An intriguing result was revealed by examining participants individually. Even for motion coherence, there was a clear suggestion of some participants showing a linear increase of MD activity across difficulty levels. These participants were matched, however, by similar numbers showing decreases. It is worth noting that, under the framework of Norman and Bobrow (1975), altered resource allocation is an option whether or not performance is actually resource-limited. When resource allocation has little effect on performance, it may vary idiosyncratically between participants.

Results were less clear for the salience of irrelevant dots. Following Han and Marois (2013), we expected that MD activity might increase with the salience of irrelevant dots, reflecting stronger attempts to focus only on the relevant red dots. Results of Experiment 1 were somewhat in line with this prediction, though even with 40 participants, the effect was not sufficiently strong to differ significantly from the null effect for coherence. In Experiment 2, even irrelevant-dots showed no hint of an overall difficulty effect. Across experiments, results for individual participants again showed a mixture of increases and decreases. Though such variable results lead to no strong interpretation, a reasonable speculation concerns variable strategies. In one extreme case, the participant could make no attempt to separate red and yellow dots, in which case yellow dots simply decrease motion coherence, and results should resemble those of a direct coherence manipulation. Under our presentation conditions, effective top-down separation of the two dot fields may have been hard or impossible to achieve, making unselective processing the dominant strategy. In some cases, however, increasing the salience of yellow dots could have increased top-down attempts to ignore them, reflected in increased MD activity.

Across many different kinds of cognitive demand, it seems that scenarios like that of Figure 1B are by far the most common. In most cases – including the rule complexity case we used in Experiment 2 – increased cognitive demand calls for increased top-down input, ensuring optimal focus on task-relevant processing. The results show, however, that this is not a universal rule. In line with the ideas of Norman and Bobrow (1975), increased attentional focus may be ineffective in overcoming simple limitations of sensory data.

As reviewed in the Introduction, the literature contains mixed findings concerning the MD response to reduced stimulus discriminability. In speeded tasks, for example, strong increases in MD activity have been reported as stimuli to be discriminated are made more similar (Jiang & Kanwisher, 2003; Woolgar et al., 2011). As we have indicated, in general we do not know the shapes of PRFs for different tasks. Rapidly deciding which of four lines is shortest, for example, may have very different attentional requirements from a global judgment of motion direction as used in the present work.

While many studies in the literature show increasing MD activity with increasing task demands, there have also been studies that have showed decreased MD activity (Bor et al., 2003), an inverted U-shape response (Callicott et al., 1999; Linden et al., 2003), or a plateau after a certain difficulty level (Marois & Ivanoff, 2005; Todd & Marois, 2004; Mitchell & Cusack, 2008). For example, Linden et al. (2003) and Callicott et al. (1999) found that the frontal-parietal network initially showed increased activation with increased working memory load, but decreased in the highest load condition close or beyond the limit of capacity. In our data there was no suggestion of an inverted-U profile; if anything, in some conditions there was a decrease in MD activity over the first few difficulty levels (e.g. Experiment 2, coherence condition). Our data suggest no decrease of MD activity as sensory limits make a task impossible to perform well.

One factor affecting MD recruitment might be advance knowledge of difficulty level. The neural basis of expectation in perceptual tasks has been shown to involve top-down modulation of frontal and parietal cortices to enhance sensory processing in the visual cortex (Giesbrecht, Weissman, Woldorff, & Mangun, 2006; Kastner, Pinsk, De Weerd, Desimone, & Ungerleider, 1999; Kok, Jehee, & de Lange, 2012; Sylvester, Shulman, Jack, & Corbetta, 2009). Furthermore, Manelis and Reder (2015) recently demonstrated using MVPA that the regions involved in a working memory task are the same regions that contain information about the upcoming task difficulty during task preparation. It is therefore possible in our study that participants in the blocked group could have decided to increase attentional effort in an attempt to compensate for anticipated perceptual difficulty. Our data, however, suggested little notable difference between mixed and blocked conditions.

Activity across the MD network is increased by many different kinds of cognitive demand (Duncan & Owen, 2000; Fedorenko et al., 2013). In contrast to this “multiple demand” pattern, the present results show little or no consistent effect of sensory manipulations in a simple motion coherence task. As suggested by Han and Marois (2013), results may reflect the degree to which performance can be improved by increasing attentional investment. When simple sensory decisions are largely data-limited, decreased accuracy cannot be compensated by increased attention, leading to little or no enhancement of MD activity.

## 5. Acknowledgements

This work was supported by funding from the Medical Research Council (United Kingdom), program SUAG/002/RG91365. TW was supported by the Taiwan Cambridge Scholarship from the Cambridge Commonwealth, European & International Trust and the Percy Lander studentship from Downing College.

## References

Baldauf, D., & Desimone, R. (2014). Neural mechanisms of object-based attention. Science, 344(6182), 424–427.

Bor, D., Duncan, J., Wiseman, R. J., & Owen, A. M. (2003). Encoding strategies dissociate prefrontal activity from working memory demand. Neuron, 37(2), 361–367.

Brett, M., Anton, J. L., Valabregue, R., & Poline, J. B. (2002). Region of interest analysis using the MarsBar toolbox for SPM 99. Neuroimage, 16(2), S497.

Callicott, J. H., Mattay, V. S., Bertolino, A., Finn, K., Coppola, R., Frank, J. A., … Weinberger, D. R. (1999). Physiological characteristics of capacity constraints in working memory as revealed by functional MRI. Cereb Cortex, 9(1), 20–26.

Chun, M. M., & Marois, R. (2002). The dark side of visual attention. Curr Opin Neurobiol, 12(2), 184–189.

Crittenden, B. M., & Duncan, J. (2014). Task difficulty manipulation reveals multiple demand activity but no frontal lobe hierarchy. Cereb Cortex, 24(2), 532–540. doi: 10.1093/cercor/bhs333

Cusack, R., Mitchell, D. J., & Duncan, J. (2010). Discrete object representation, attention switching, and task difficulty in the parietal lobe. J Cogn Neurosci, 22(1), 32–47. doi: 10.1162/jocn.2009.21194

Cusack, R., Vicente-Grabovetsky, A., Mitchell, D. J., Wild, C. J., Auer, T., Linke, A. C., & Peelle, J. E. (2014). Automatic analysis (aa): efficient neuroimaging workflows and parallel processing using Matlab and XML. Front Neuroinform, 8, 90. doi: 10.3389/fninf.2014.00090

Deary, I. J., Simonotto, E., Meyer, M., Marshall, A., Marshall, I., Goddard, N., & Wardlaw, J. M. (2004). The functional anatomy of inspection time: an event-related fMRI study. Neuroimage, 22(4), 1466–1479.

Desimone, R., & Duncan, J. (1995). Neural mechanisms of selective visual attention. Annu Rev Neurosci, 18, 193–222. doi: 10.1146/annurev.ne.18.030195.001205

Duncan, J. (2010). The multiple-demand (MD) system of the primate brain: mental programs for intelligent behaviour. Trends Cogn Sci, 14(4), 172–179. doi: 10.1016/j.tics.2010.01.004

Duncan, J., & Owen, A. M. (2000). Common regions of the human frontal lobe recruited by diverse cognitive demands. Trends in Neurosciences, 23(10), 475–483.

Farooqui, A. A., Mitchell, D., Thompson, R., & Duncan, J. (2012). Hierarchical organization of cognition reflected in distributed frontoparietal activity. Journal of Neuroscience, 32(48), 17373–17381.

Fedorenko, E., Duncan, J., & Kanwisher, N. (2013). Broad domain generality in focal regions of frontal and parietal cortex. Proc Natl Acad Sci U S A, 110(41), 16616–16621. doi: 10.1073/pnas.1315235110

Fox, M. D., Snyder, A. Z., Vincent, J. L., Corbetta, M., Van Essen, D. C., & Raichle, M. E. (2005). The human brain is intrinsically organized into dynamic, anticorrelated functional networks. Proc Natl Acad Sci U S A, 102(27), 9673–9678. doi: 10.1073/pnas.0504136102

Giesbrecht, B., Weissman, D. H., Woldorff, M. G., & Mangun, G. R. (2006). Pre-target activity in visual cortex predicts behavioral performance on spatial and feature attention tasks. Brain Res, 1080(1), 63–72. doi: 10.1016/j.brainres.2005.09.068

Han, S. W., & Marois, R. (2013). Dissociation between process-based and data-based limitations for conscious perception in the human brain. Neuroimage, 64, 399–406. doi: 10.1016/j.neuroimage.2012.09.016

Heekeren, H. R., Marrett, S., Bandettini, P. A., & Ungerleider, L. G. (2004). A general mechanism for perceptual decision-making in the human brain. Nature, 431(7010), 859–862. doi: 10.1038/nature02966

Heekeren, H. R., Marrett, S., Ruff, D. A., Bandettini, P. A., & Ungerleider, L. G. (2006). Involvement of human left dorsolateral prefrontal cortex in perceptual decision making is independent of response modality. Proc Natl Acad Sci U S A, 103(26), 10023–10028. doi: 10.1073/pnas.0603949103

Heekeren, H. R., Marrett, S., & Ungerleider, L. G. (2008). The neural systems that mediate human perceptual decision making. Nat Rev Neurosci, 9(6), 467–479. doi: 10.1038/nrn2374

Heller, R., Golland, Y., Malach, R., & Benjamini, Y. (2007). Conjunction group analysis: an alternative to mixed/random effect analysis. Neuroimage, 37(4), 1178–1185.

Holcomb, H. H., Medoff, D. R., Caudill, P. J., Zhao, Z., Lahti, A. C., Dannals, R. F., & Tamminga, C. A. (1998). Cerebral blood flow relationships associated with a difficult tone recognition task in trained normal volunteers. Cerebral Cortex, 8(6), 534–542.

Hou, Y., & Liu, T. (2012). Neural correlates of object-based attentional selection in human cortex. Neuropsychologia, 50(12), 2916–2925. doi: 10.1016/j.neuropsychologia.2012.08.022

Jiang, Y., & Kanwisher, N. (2003). Common neural mechanisms for response selection and perceptual processing. Journal of Cognitive Neuroscience, 15(8), 1095–1110.

Jovicich, J., Peters, R. J., Koch, C., Braun, J., Chang, L., & Ernst, T. (2001). Brain areas specific for attentional load in a motion-tracking task. J Cogn Neurosci, 13(8), 1048–1058. doi: 10.1162/089892901753294347

Kastner, S., Pinsk, M. A., De Weerd, P., Desimone, R., & Ungerleider, L. G. (1999). Increased activity in human visual cortex during directed attention in the absence of visual stimulation. Neuron, 22(4), 751–761.

Kayser, A. S., Buchsbaum, B. R., Erickson, D. T., & D’Esposito, M. (2010). The functional anatomy of a perceptual decision in the human brain. Journal of Neurophysiology, 22 103(3), 1179–1194. doi: 10.1152/jn.00364.2009

Kayser, A. S., Erickson, D. T., Buchsbaum, B. R., & D’Esposito, M. (2010). Neural representations of relevant and irrelevant features in perceptual decision making. J Neurosci, 30(47), 15778–15789. doi: 10.1523/JNEUROSCI.3163-10.2010

Kiehl, K. A., Liddle, P. F., & Hopfinger, J. B. (2000). Error processing and the rostral anterior cingulate: An event□related fMRI study. Psychophysiology, 37(2), 216–223.

Kim, J. N., & Shadlen, M. N. (1999). Neural correlates of a decision in the dorsolateral prefrontal cortex of the macaque. Nature Neuroscience, 2(2), 176–185. doi: 10.1038/5739

Kok, P., Jehee, J. F., & de Lange, F. P. (2012). Less is more: expectation sharpens representations in the primary visual cortex. Neuron, 75(2), 265–270. doi: 10.1016/j.neuron.2012.04.034

Linden, D. E., Bittner, R. A., Muckli, L., Waltz, J. A., Kriegeskorte, N., Goebel, R., … Munk, M. H. (2003). Cortical capacity constraints for visual working memory: dissociation of fMRI load effects in a fronto-parietal network. Neuroimage, 20(3), 1518–1530.

Manelis, A., & Reder, L. M. (2015). He who is well prepared has half won the battle: an FMRI study of task preparation. Cereb Cortex, 25(3), 726–735. doi: 10.1093/cercor/bht262

Marois, R., Chun, M. M., & Gore, J. C. (2004). A common parieto-frontal network is recruited under both low visibility and high perceptual interference conditions. Journal of Neurophysiology, 92(5), 2985–2992. doi: 10.1152/jn.01061.2003

Marois, R., & Ivanoff, J. (2005). Capacity limits of information processing in the brain. Trends Cogn Sci, 9(6), 296–305. doi: 10.1016/j.tics.2005.04.010

Martinez-Trujillo, J., & Treue, S. (2002). Attentional modulation strength in cortical area MT depends on stimulus contrast. Neuron, 35(2), 365–370.

Miller, E. K., & Cohen, J. D. (2001). An integrative theory of prefrontal cortex function. Annual review of neuroscience, 24(1), 167–202.

Mitchell, D. J., & Cusack, R. (2008). Flexible, capacity-limited activity of posterior parietal cortex in perceptual as well as visual short-term memory tasks. Cerebral Cortex, 18(8), 1788–1798.

Muller-Gass, A., & Schroger, E. (2007). Perceptual and cognitive task difficulty has differential effects on auditory distraction. Brain Res, 1136(1), 169–177. doi: 10.1016/j.brainres.2006.12.020

Niendam, T. A., Laird, A. R., Ray, K. L., Dean, Y. M., Glahn, D. C., & Carter, C. S. (2012). Meta-analytic evidence for a superordinate cognitive control network subserving diverse executive functions. Cogn Affect Behav Neurosci, 12(2), 241–268. doi: 10.3758/s13415-011-0083-5

Norman, D. A., & Bobrow, D. G. (1975). On data-limited and resource-limited processes. Cognitive psychology, 7(1), 44–64.

Norman, D. A., & Shallice, T. (1980). Attention to action: Willed and automatic control of behavior (No. CHIP-99). CALIFORNIA UNIV SAN DIEGO LA JOLLA CENTER FOR HUMAN INFORMATION PROCESSING.

Rees, G., Friston, K., & Koch, C. (2000). A direct quantitative relationship between the functional properties of human and macaque V5. Nature Neuroscience, 3(7), 716–723. doi: 10.1038/76673

Reynolds, J. H., Pasternak, T., & Desimone, R. (2000). Attention increases sensitivity of V4 neurons. Neuron, 26(3), 703–714.

Rothman, K. J. (1990). No adjustments are needed for multiple comparisons. Epidemiology, 1(1), 43–46.

Saville, D. J. (2003). Basic statistics and the inconsistency of multiple comparison procedures. Can J Exp Psychol, 57(3), 167–175.

Seidl, K. N., Peelen, M. V., & Kastner, S. (2012). Neural evidence for distracter suppression during visual search in real-world scenes. J Neurosci, 32(34), 11812–11819. doi: 10.1523/JNEUROSCI.1693-12.2012

Spitzer, H., Desimone, R., & Moran, J. (1988). Increased attention enhances both behavioral and neuronal performance. Science, 240(4850), 338–340.

Stiers, P., Mennes, M., & Sunaert, S. (2010). Distributed task coding throughout the multiple demand network of the human frontal-insular cortex. Neuroimage, 52(1), 252–262. doi: 10.1016/j.neuroimage.2010.03.078

Sunaert, S., Van Hecke, P., Marchal, G., & Orban, G. A. (2000). Attention to speed of motion, speed discrimination, and task difficulty: an fMRI study. Neuroimage, 11(6 Pt 1), 612–623. doi: 10.1006/nimg.2000.0587

Sylvester, C. M., Shulman, G. L., Jack, A. I., & Corbetta, M. (2009). Anticipatory and stimulus-evoked blood oxygenation level-dependent modulations related to spatial attention reflect a common additive signal. J Neurosci, 29(34), 10671–10682. doi: 10.1523/JNEUROSCI.1141-09.2009

Todd, J. J., & Marois, R. (2004). Capacity limit of visual short-term memory in human posterior parietal cortex. Nature, 428(6984), 751–754. doi: 10.1038/nature02466

Ullsperger, M., & Von Cramon, D. Y. (2004). Neuroimaging of performance monitoring: error detection and beyond. Cortex, 40(4), 593–604.

Wager, T. D., Jonides, J., & Reading, S. (2004). Neuroimaging studies of shifting attention: a meta-analysis. Neuroimage, 22(4), 1679–1693.

Woolgar, A., Afshar, S., Williams, M. A., & Rich, A. N. (2015). Flexible Coding of Task Rules in Frontoparietal Cortex: An Adaptive System for Flexible Cognitive Control. J Cogn Neurosci, 27(10), 1895–1911. doi: 10.1162/jocn_a_00827

Woolgar, A., Hampshire, A., Thompson, R., & Duncan, J. (2011). Adaptive coding of task-relevant information in human frontoparietal cortex. J Neurosci, 31(41), 14592–14599. doi: 10.1523/JNEUROSCI.2616-11.2011

Woolgar, A., Williams, M. A., & Rich, A. N. (2015). Attention enhances multi-voxel representation of novel objects in frontal, parietal and visual cortices. Neuroimage, 109, 429–437. doi: 10.1016/j.neuroimage.2014.12.083

